# Early conservation benefits of a de facto marine protected area at San Clemente Island, California

**DOI:** 10.1101/796193

**Authors:** Michael W. Esgro, James Lindholm, Kerry J. Nickols, Jessica Bredvik

**Affiliations:** Institute for Applied Marine Ecology, California State University Monterey Bay, Seaside, CA 93955; California State University Northridge, Northridge, CA 91330; Naval Facilities Engineering Command Southwest, San Diego, CA 92132

## Abstract

De facto marine protected areas (DFMPAs) are regions of the ocean where human activity is restricted for reasons other than conservation. Although DFMPAs are widespread globally, their potential role in the protection of marine habitats, species, and ecosystems has not been well studied. In 2012 and 2013, we conducted remotely operated vehicle (ROV) surveys of marine communities at a military DFMPA and an adjacent fished reference site at San Clemente Island, the southernmost of California’s Channel Islands. We used data extracted from ROV imagery to compare density and biomass of focal species, as well as biodiversity and community composition, between the two sites. Generalized linear modeling indicated that both density and biomass of California sheephead (*Semicossyphus pulcher*) were significantly higher inside the DFMPA. Biomass of ocean whitefish (*Caulolatilus princeps*) was also significantly higher inside the DFMPA. However, species richness and Shannon-Weaver diversity were not significantly higher inside the DFMPA, and overall fish community composition did not differ significantly between sites. Demonstrable differences between the DFMPA and fished site for two highly sought-after species hint at early potential benefits of protection, though the lack of differences in the broader community suggests that a longer trajectory of recovery may be required for other species. A more comprehensive understanding of the potential conservation benefits of DFMPAs is important in the context of marine spatial planning and global marine conservation objectives.

## Introduction

Marine ecosystems worldwide are threatened by a variety of stressors, including overharvest, pollution, and climate change [1–4]. To effectively manage these complex and often interrelated problems, policymakers are increasingly adopting marine spatial planning (MSP) as a management technique [5–8]. MSP is an integrative, ecosystem-based framework that accounts for the effects of multiple human uses on marine systems and informs the spatial distribution of these activities. When used effectively and in concert with other marine management tools, MSP can safeguard ocean health while maintaining the delivery of essential ecosystem services [7].

Marine protected areas (MPAs), regions of the ocean set aside for conservation, are key components of MSP [6–7]. The number of MPAs has increased dramatically in recent years, with the percentage of the global ocean that is “strongly” or “fully” protected increasing from less than 0.1% to 0.6% in the last decade [9–11]. The realized ecological and economic benefits of MPAs vary widely depending on level of protection [11]. However, there is general agreement among the scientific community that strongly or fully protected MPAs can increase species density and biomass, promote the recovery of size-truncated populations, and increase biodiversity within and beyond their boundaries [9, 12].

Despite the preponderance of evidence for the conservation benefits of MPAs, very little research has examined the potentially similar benefits of de facto marine protected areas (DFMPAs)—places where human activity is restricted by law for reasons other than conservation or natural resource management [13]. Examples of DFMPAs include restricted areas reserved for military use [13], cable exclusion zones [14], and marine renewable energy installations such as wind turbines [15]. The first comprehensive inventory of DFMPAs in the United States indicated that there were more than 1,200 DFMPAs within U.S. waters, covering an area roughly equivalent to the total combined area protected by state and federal MPAs [16].

DFMPAs likely play a critical and heretofore unappreciated role in marine conservation [16]. On land, restricted areas such as military bases have been shown to contain higher densities of threatened and endangered species, as well as higher overall biodiversity, when compared to adjacent areas open to public access [17]. In the marine environment, Roberts et al. (2001) analyzed catch data from several Florida coast fisheries and found significantly higher numbers of world-record sized catches in fisheries located near the Merritt Island National Wildlife Refuge, access to which is restricted due to the Refuge’s proximity to NASA’s Kennedy Space Center – potential evidence of spillover from a DFMPA [18]. Rogers-Bennett et al. (2013) compared historical intertidal red abalone (*Haliotis rufescens*) densities inside and outside a DFMPA (the Stornetta Ranch property) on California’s central coast, which was closed to the public from 1917-2004. The authors documented 86% higher abalone densities inside the DFMPA compared to adjacent areas before the area was opened to fishing in 2004 [19].

A more comprehensive understanding of how DFMPAs contribute to marine conservation is essential in the context of MSP. In California, for example, the Marine Life Protection Act requires that the state’s system of MPAs be designed and managed as an ecologically cohesive network [20, 21]. It is likely that DFMPAs make nontrivial ecological contributions to that network, for example as sources or sinks of larval organisms, but the paucity of information regarding DFMPAs has largely precluded their incorporation into California’s MPA management efforts to date.

San Clemente Island (SCI), the southernmost of the Channel Islands in the Southern California Bight (Fig 1), has been owned and managed by the United States Navy since 1934. SCI supports vital military activities that cannot be conducted anywhere else in the world [22]. SCI’s waters are also home to highly productive and economically important fisheries, both commercial and recreational. Civilians regularly use the waters surrounding SCI for non-consumptive recreational activities such as boating and SCUBA diving [22, 23]. However, civilian access to areas in which certain military training exercises are conducted is highly restricted and in some cases prohibited. To safely facilitate multiple human uses at SCI, the waters surrounding the island up to 3 nautical miles have been divided into eight naval safety zones [24] (Fig 1). The type and frequency of military use, as well as associated restrictions on civilian access and activity, differ from zone to zone. Two locations, Zone G and Wilson Cove, are permanently closed to civilians. Other zones are only closed when being used for military activities that pose a threat to public safety [24]. The presence of both restricted and unrestricted areas at SCI presents a unique opportunity to compare DFMPAs to fished areas of similar habitat quality and habitat type distribution.

**Fig 1.**
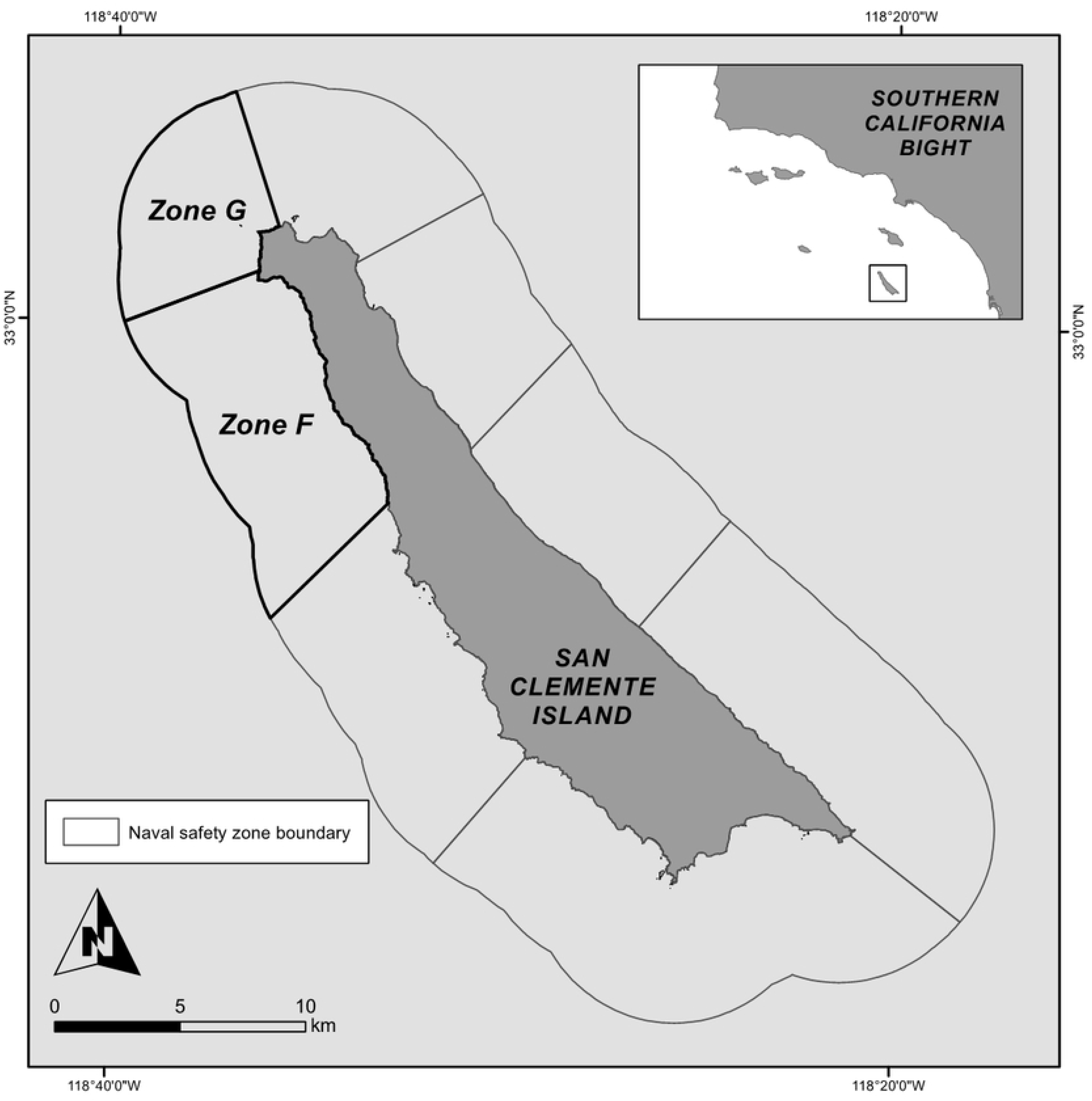
San Clemente Island. San Clemente Island is the southernmost of the Channel Islands in the California Bight. To safely facilitate multiple uses at San Clemente Island, the surrounding waters up to 3 nautical miles have been divided into eight naval safety zones. This study compared marine communities at two naval safety zones at the northwest corner of the island: a DFMPA site (Zone G) and an adjacent fished reference site (Zone F).

We used remotely operated vehicle (ROV) imagery to compare marine communities at a DFMPA site and an adjacent fished reference site at SCI. Specifically, we tested the following hypotheses: (1) density of fished species is higher at the DFMPA site than at the fished site, (2) biomass of fished species is higher at the DFMPA site than at the fished site, (3) species richness and Shannon-Weaver diversity are higher at the DFMPA site than at the fished site, and (4) fish community composition differs between sites. Our objective was to assess the potential conservation benefits of the DFMPA using these well-established MPA performance indicators.

## Materials and Methods

### Study site

SCI is located 70 km west of the U.S. mainland and 30 km south of Santa Catalina Island in the Southern California Bight, and is home to a diverse assemblage of marine flora and fauna [25, 26, 27]. We compared marine communities at two sites at northwest SCI: Naval Safety Zone G (118°38’3.259” W, 33°2’1.831” N) and Naval Safety Zone F (118°36’8.296” W, 32°59’27.276” N) (Fig 1). Zone G is used regularly for Navy SEAL training, live-fire practice, and other military activity; it has been closed to all civilian access since June 2010 [24]. Zone F is open to civilians except for occasional short closures when military activities are being conducted that might threaten public safety; it is commonly fished by recreational anglers [24].

### Image collection

ROV imagery was collected over the course of two week-long cruises in November 2012 and 2013 using a Vector M4 ROV (Deep Ocean Engineering, San Jose, California) deployed from a fishing vessel. ROV configuration and sampling protocols were based on previous studies conducted by the authors and collaborators [28, 29, 30]. The ROV was equipped with five cameras (forward-facing standard-definition video, forward-facing high-definition video, down-facing standard-definition video, digital high-definition still, and rear facing safety video), halogen lights, paired forward- and down-facing sizing lasers spaced 10 cm apart, a strobe for still photos, an altimeter, and forward-facing multibeam sonar. While at depth, the position of the ROV on the seafloor was maintained by a Trackpoint III acoustic positioning system, with the resulting coordinates logged into Hypack navigational software.

The ROV was flown over the seafloor along predetermined transect lines at a mean altitude of 1.0 m and a speed of approximately 0.67 knots. Transect placement was designed to sample a variety of depths and habitats and was based on *a priori* analysis of existing seafloor mapping data (Fig 2). While on transect, continuous video imagery was recorded from the ROV’s cameras to digital tape. Still images were collected opportunistically along each transect.

**Fig 2.**
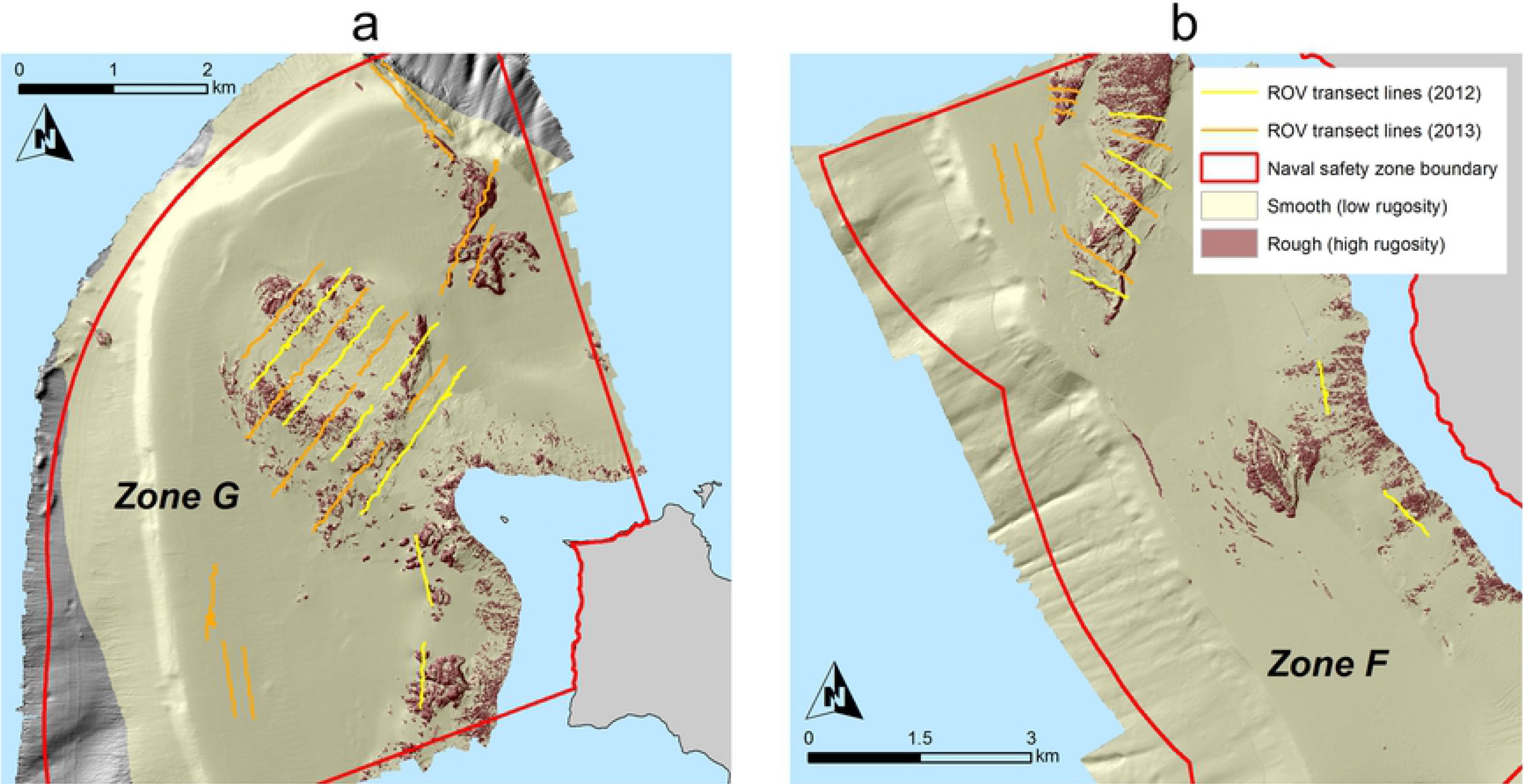
Seafloor maps of a) the DFMPA site (Zone G) and b) the fished site (Zone F). High rugosity areas indicate rocky substrate; low rugosity areas indicate sandy substrate. Transect placement was designed to encompass a variety of depths and habitat types.

### Focal species

We compared density and biomass between the two sites for focal species associated with a range of habitats and trophic levels (Table 1). We selected focal species that met the following criteria: (1) targeted for long-term monitoring in California due to ecological and/or economic importance, (2) sufficiently abundant at SCI in 2012 and 2013 to allow for reasonable sample sizes, and (3) easily identifiable in ROV video and photo imagery.

**Table 1.** Focal species list and categorization.

| Rock-associated focal species |
| --- |
| Predatory fishes |
| Lingcod (*Ophiodon elongatus*) |
| California sheephead (*Semicossyphus pulcher*) |
| California scorpionfish (*Scorpaena guttata*) |
| Ocean whitefish (*Caulolatilus princeps*) |
| Bocaccio rockfish (*Sebastes paucispinis*) |
| Copper rockfish (*Sebastes caurinus*) |
| Olive/yellowtail rockfish (*Sebastes serranoides*/*S. flavidus*) |
| Vermilion/canary rockfish (*Sebastes miniatus*/*S. pinniger*) |
| Dwarf rockfishes |
| Dwarf-red rockfish (*Sebastes rufianus*) |
| Halfbanded rockfish (*Sebastes semicinctus*) |
| Squarespot rockfish (*Sebastes hopkinsi*) |
| Mobile invertebrates |
| California spiny lobster (*Panulirus interruptus*) |
| Sand-associated focal species |
| Predatory fishes |
| Sanddab (*Citharichthys* spp.) |
| Surfperch (Embiotocidae, multiple species) |
| Mobile invertebrates |
| California sea cucumber (*Parastichopus californicus*) |

This study focused on mid-depth (40-200 m) demersal communities, as mid-depth rock habitat represents at least 75% of all marine habitats in California state waters by area and supports a high diversity of ecologically and economically important demersal fish and invertebrate species – many of which have been listed by the California Department of Fish and Wildlife as “likely to benefit” from MPA protection [31].

### Data extraction from imagery

All video imagery was reviewed as a series of non-overlapping “video quadrats.” For each individual organism encountered in forward facing-video we noted the time of occurrence and identified the organism to the lowest taxonomic level possible. Identification was aided by still images and downward-facing video. Organism sizes (total lengths) were estimated to the nearest 5 cm using the paired sizing lasers and grouped into 5 cm size bins. For fishes, these lengths were later converted to weights (kg) using size class midpoints and the length-weight relationship (LWR)

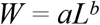

where *W* is weight of a fish in kg, *L* is the length of that fish in cm, and *a* and *b* are constants unique to individual fish species (Table S1).

Physical habitat data were collected separately by re-watching each ROV transect. Organisms were ignored during this round of data collection, and the video was paused to record the dominant habitat in each non-overlapping quadrat (> 50% of the quadrat) as rock, sand, or mixed.

### Analysis

Percent rock was calculated based on analysis of video-derived physical habitat data, and represented the percentage of video quadrats on each transect that were classified as rock or mixed habitat. Mean transect depths were calculated from data generated by the ROV’s navigational sensors, which recorded depth every second while the ROV was on transect. In general, ROV transects closely followed bathymetric contours, so depth did not vary substantially over the course of transects. Area surveyed was calculated by multiplying transect length by transect width, assumed to be 1 m for all transects based on the field of view of the ROV’s cameras. In our analyses, described in more detail below, we considered these variables along with site (DFMPA or fished) as possible predictors of density, biomass, richness, and diversity, with individual transects serving as the unit of replication. Sampling year was not included as a predictor variable in our analyses, as this study was not designed for temporal comparison, i.e. transects were not resampled in the second sampling year. Values for these variables associated with each transect are reported in Table S2.

To analyze potential relatedness between predictor variables, we conducted a Factor Analysis of Mixed Data (FAMD) using the package FactoMineR in R [32]. FAMD is designed to analyze relationships among both continuous and categorical variables; it functions as a Principal Components Analysis for continuous data (mean depth, percent rock, area surveyed) and a Multiple Correspondence Analysis for categorical data (site). Squared correlation coefficients determined degree of relatedness between variables. We also compared mean percent rock and mean depth between control and DFMPA transects using one-way analysis of variance (ANOVA).

For species-level comparisons, density was calculated by dividing total number of organisms on a given transect by total area surveyed during the transect. Biomass was calculated by dividing total weight of organisms on a given transect by total area surveyed during the transect. For community-level comparisons, standardized species richness was calculated by dividing the total number of unique species on a given transect by the area surveyed during the transect. Standardized species diversity was calculated by first computing the Shannon-Weaver diversity index on a given transect, and then dividing by the area surveyed during the transect.

Mean density, biomass, richness, and Shannon-Weaver diversity were assessed using the following generalized linear model with a log link function:

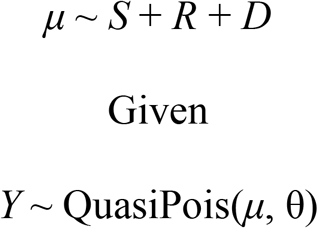

where *Y* is a random variable representing the ecological metric of interest, quasi-Poisson distributed with mean *μ* and variance θ; *S* is a categorical variable representing site, *R* is a continuous variable representing percent rock, and *D* is a continuous variable representing depth. We compared models containing all possible combinations of predictor variables, including a null model, using Akaike’s Information Criterion (AIC).

To explore differences in fish community composition between sites, we calculated Bray-Curtis dissimilarity indices between all possible transect pairs. The Bray-Curtis dissimilarity index [33] quantifies the dissimilarity in species composition between two sites based on counts per area of unique fish species at each site. These calculations were based on all unique fish species observed along transects, not just focal species. Bray-Curtis dissimilarity indices were used to conduct an analysis of similarity for fish communities between sites.

All statistical analysis was conducted using R statistical software and associated packages, version 2.14.1 [34].

## Results

We conducted 15 transects (13,593.39 m^2^ surveyed) at the fished site and 19 transects (25,264.01 m^2^ surveyed) at the DFMPA site (Fig 2). A total of 51,688 fishes, representing 64 distinct species or species groups, and 184 mobile invertebrates, representing 8 distinct species or species groups, were observed. We also encountered a wide variety of sessile invertebrates including corals, sponges, sea whips, and sea pens in both years at both sites.

Factor analysis of mixed data indicated that site was correlated with sampling effort (i.e. more sampling occurred in the DFMPA) and percent rock was correlated with depth. However, neither percent rock nor depth was correlated with site, indicating that there were no significant differences in mean depth or mean percent rock between sites. This was confirmed by statistical comparison of mean percent rock between fished site and DFMPA site transects (one-way ANOVA, F = 0.12, p = 0.73), and mean depth between fished site and DFMPA site transects (one-way ANOVA, F = 0, p = 0.99).

Mean density and biomass for all focal species are shown in Fig 3 and reported in Tables S3-S4. Site was found to be a significant predictor of increased California sheephead density, California sheephead biomass, and ocean whitefish biomass, with all of these metrics higher at the DFMPA than at the fished site. For most other focal species, percent rock and/or depth were the only significant predictors of density and biomass. For some species, no variables were found to be significant predictors of density or biomass (i.e. for these species the null model had the lowest AIC value). For density, these species were: California scorpionfish, olive/yellowtail rockfish, dwarf-red rockfish, squarespot rockfish, surfperch, and sea cucumbers. For biomass, these species were: California scorpionfish, copper rockfish, squarespot rockfish, and surfperch (Tables 2-3).

**Fig 3.**
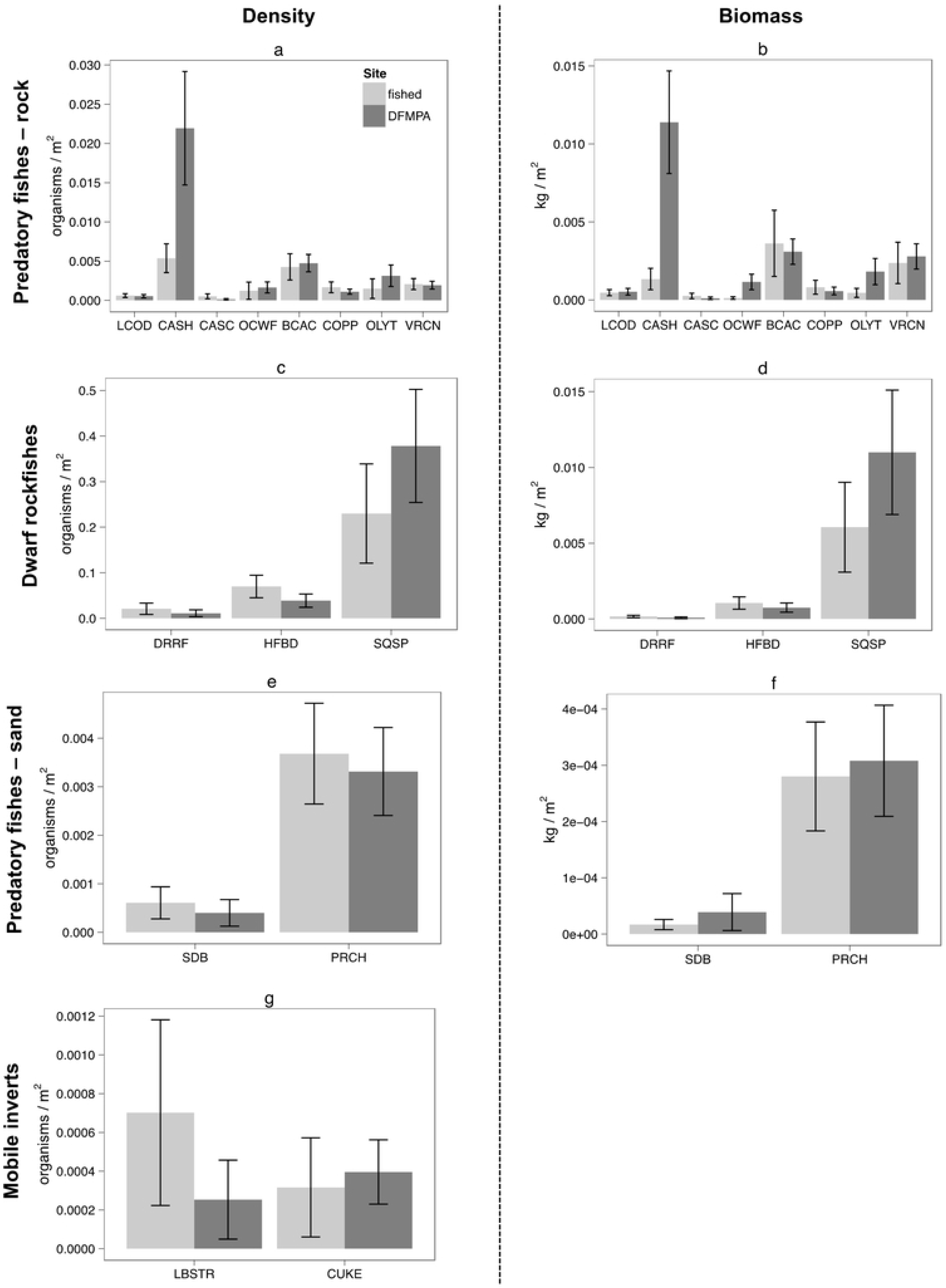
Mean density and biomass for all focal species. a) Density comparisons for rock-associated predatory fishes, b) Biomass comparisons for rock-associated predatory fishes, c) Density comparisons for dwarf rockfishes, d) Biomass comparisons for dwarf rockfishes, e) Density comparisons for sand-associated predatory fishes, f) Biomass comparisons for sand-associated predatory fishes, g) Density comparisons for mobile invertebrates. Abbreviations: LCOD = lingcod, CASH = California sheephead, CASC = California scorpionfish, OCWF = ocean whitefish, BCAC = bocaccio rockfish, COPP = copper rockfish, OLYT = olive/yellowtail rockfish, VRCN = vermilion/canary rockfish, DRRF = dwarf-red rockfish, HFBD = halfbanded rockfish, SQSP = squarespot rockfish, SDB = sanddab, PRCH = perch, LBSTR = California spiny lobster, CUKE = California sea cucumber.

**Table 2.**
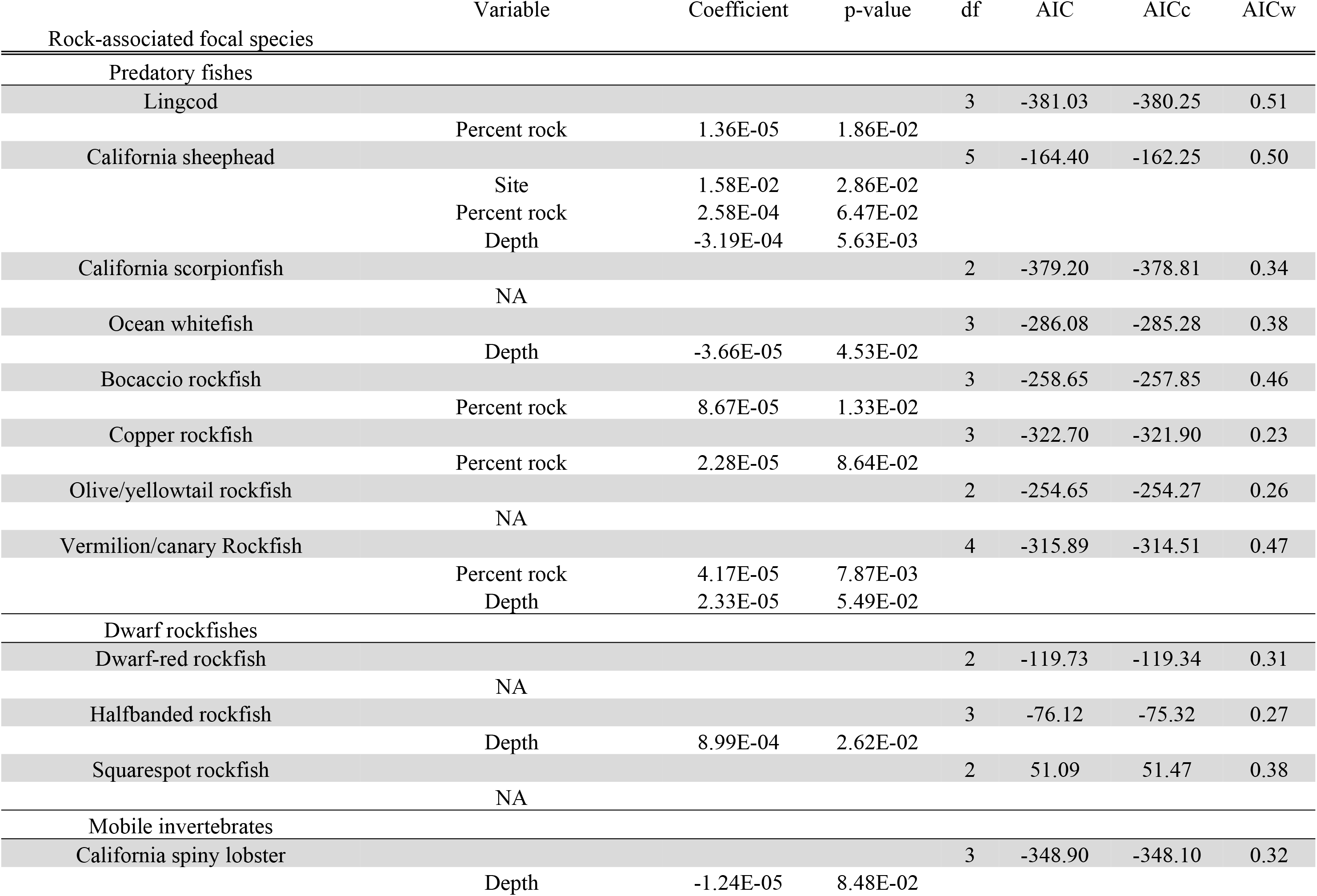
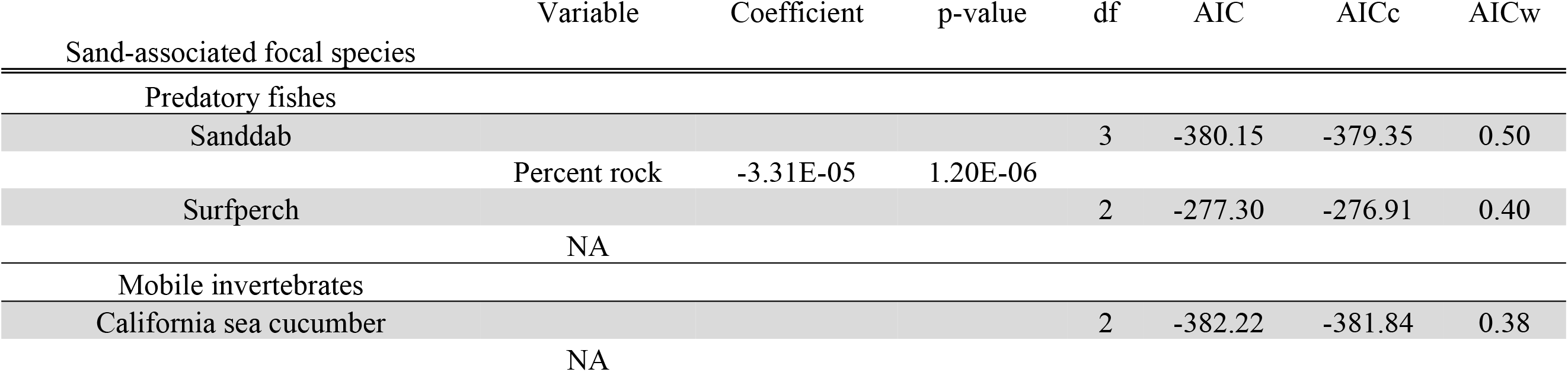
GLM results for focal species density. Models shown are those with the lowest AICc of all candidate models considered.

**Table 3.**
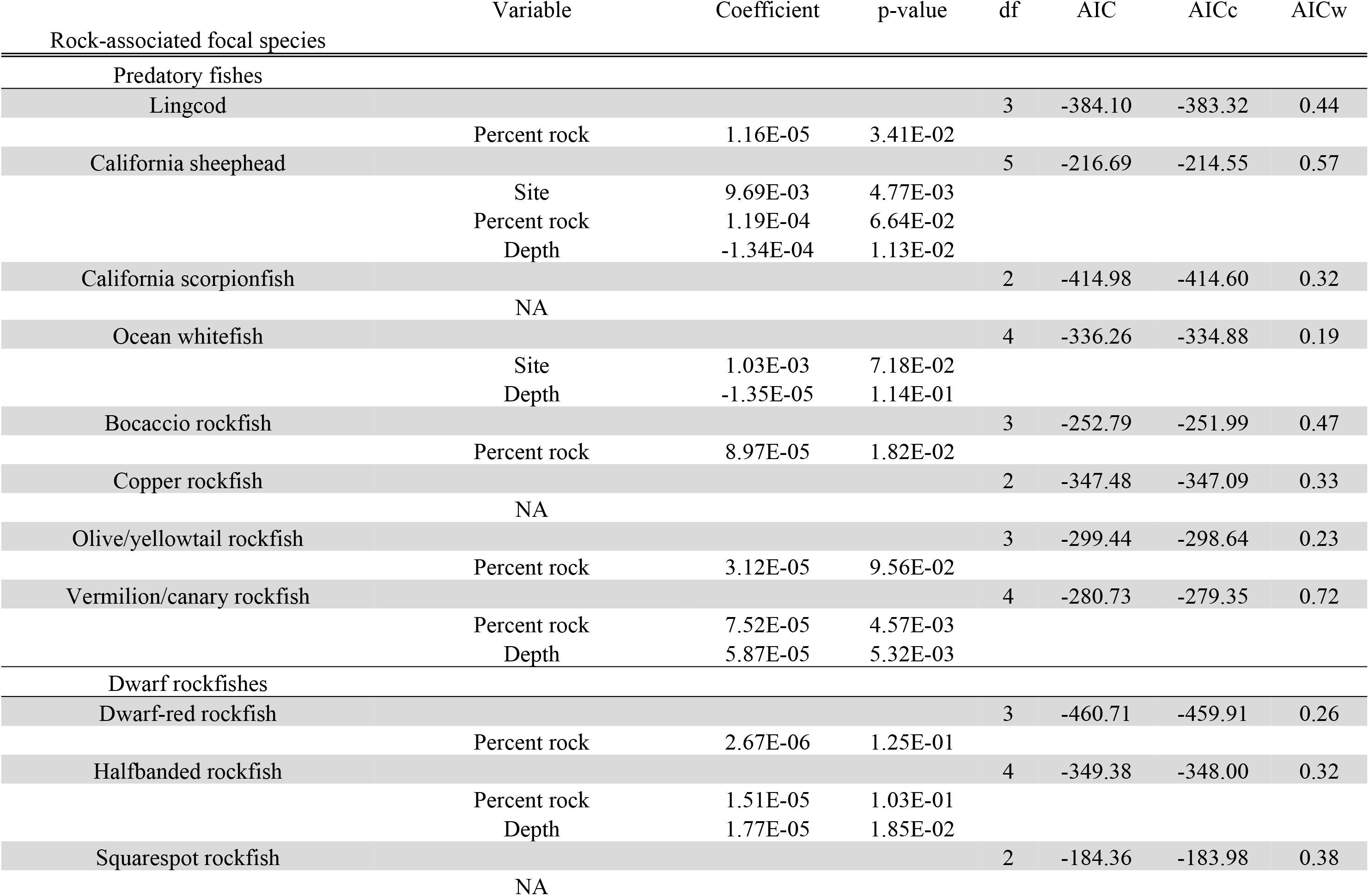
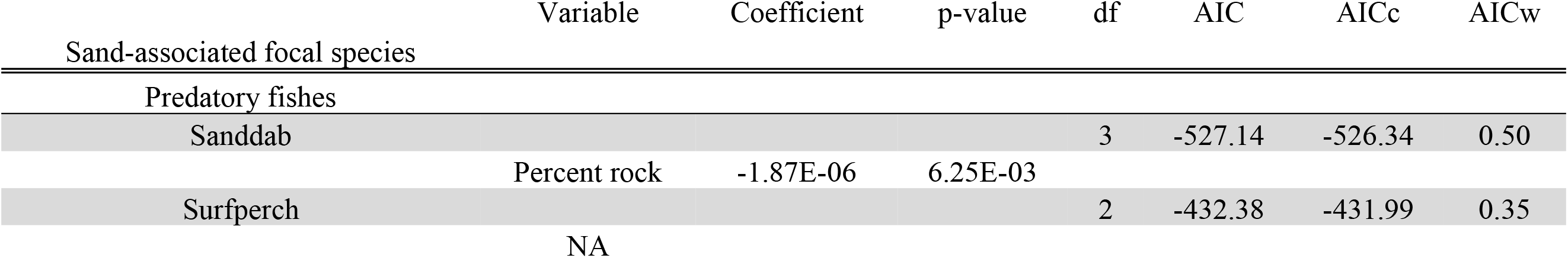
GLM results for focal species biomass. Models shown are those with the lowest AICc of all candidate models considered.

Site and percent rock were significant predictors of species richness, with richness significantly lower at the DFMPA site, while only depth was a significant predictor of Shannon-Weaver diversity (Table 4). Non-metric multidimensional scaling based on Bray-Curtis dissimilarity indices between transects and an analysis of similarity showed no significant differences in fish communities between sites (R = 0.031, p = 0.18).

**Table 4.** GLM results for mean standardized species richness and mean standardized Shannon-Weaver diversity. Models shown are those with the lowest AICc of all candidate models considered.

|  | Variable | Coefficient | p-value | df | AIC | AICc | AICw |
| --- | --- | --- | --- | --- | --- | --- | --- |
| Richness |  |  |  | 4 | -234.69 | -233.44 | 0.63 |
|  | Site | -6.20E-03 | 1.82E-02 |  |  |  |  |
|  | Percent rock | 2.00E-04 | 1.00E-04 |  |  |  |  |
| Diversity |  |  |  | 3 | -382.10 | -381.30 | 0.24 |
|  | Depth | -6.68E-06 | 1.29E-01 |  |  |  |  |

## Discussion

Demonstrable differences between the DFMPA and fished sites for two highly sought-after fishes, California sheephead and ocean whitefish, hint at hint at early potential benefits of protection, though the lack of differences in the broader community suggests a longer trajectory of recovery may be required for other species. We hypothesized that reduced fishing pressure in the DFMPA might result in conservation benefits similar to those that have been documented in MPAs across the globe — namely, increased density and biomass of some fished species, as well as community-wide effects including increased species richness and diversity [9]. However, we recognized that the relatively short two to three-year period of “recovery” at the time of the ROV surveys may limit those observed benefits [35–36]. Indeed, we observed very limited evidence for our hypotheses, with only 1 of 15 focal species showing increases in density and 2 of 15 focal species showing increases in biomass at the DFMPA site.

California sheephead exhibited the most striking result – ten-fold increases in both density and biomass at the DFMPA site. California sheephead are highly sought after by recreational anglers in Southern California. Large males in particular are preferred targets for fishermen [37–38]; since large males monopolize access to female sheephead, their removal can dramatically reduce the species’ overall reproductive rate in fished areas [37, 39]. This effect is compounded by the fact that sheephead are protogynous sequential hermaphrodites, which means that in the absence of a male reproduction is halted until a female can transition sexes and take its place [39]. These life history characteristics, coupled with high historical fishing pressure in Southern California, make sheephead a potential bellwether for ecological changes resulting from the reduction or removal of fishing pressure.

Biomass of ocean whitefish was significantly higher at the DFMPA site. Like sheephead, ocean whitefish are commonly fished by recreational anglers in Southern California. Increases in mean body size and filling in of size-truncated populations is a well-documented response to protection [9, 35–36]. We explored this question further by comparing size frequency distributions for all species. California sheephead did indeed show potential filling in of size classes (Fig 4), but the distributions were not significantly different between fished and DFMPA sites according to Kolmogorov-Smirnov tests (p = 0.07). We did not find a positive relationship between protection and biomass, or any evidence of changes in size frequency distribution, for any of the other species considered.

**Fig 4.**
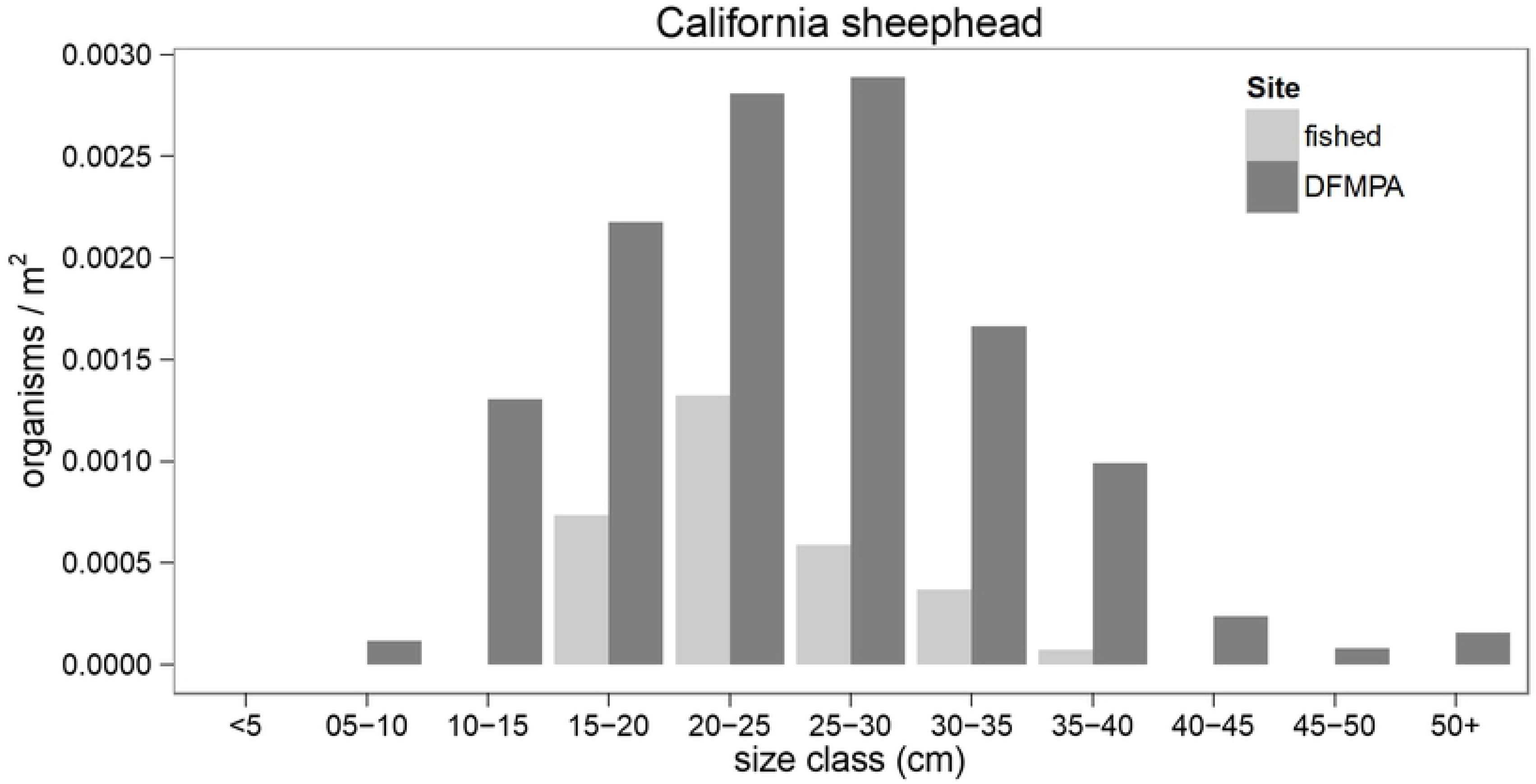
California sheephead size frequency distribution. Comparison of California sheephead size frequencies at the DFMPA and fished sites.

The lack of observable community-wide differences between sites may have been due, at least in part, to differences in habitat quality, the spillover effect, or other human uses. Habitat/microhabitat type, quality, and availability are critical drivers of marine species distribution and community composition, and in some cases are more influential than the presence or absence of protection from fishing [40–43]. In addition, physical and chemical oceanographic conditions have significant impacts on marine communities, for example by driving patterns of larval dispersal or influencing nutrient availability in an ecosystem [44–46]. These factors have the potential to override or confound any potential benefits of removing or reducing fishing pressure. However, the results of our habitat analyses suggested that the DFMPA and fished sites were similar in terms of habitat, an important consideration given the fact that depth and percent rock were found to be significant predictors of many of the ecological metrics we examined. Further study using advanced habitat suitability modeling techniques and physical oceanographic modeling would allow for a more fine-scale comparison of the distribution of suitable habitat for focal species between sites [30, 47].

A lack of ecological divergence between sites may also have been a result of the spillover effect. Spillover of adult or larval organisms from MPAs to unprotected waters is widely acknowledged as an economic benefit of spatial protection [18, 48–49]. However, spillover may confound spatial comparison if organisms are exported from a protected site to a reference site. This confounding factor is especially important to consider when the protected and control sites are close together [50], as was the case for the sites in this study. Including more of SCI’s closed and fished sites in a larger-scale analysis, with sites serving as units of replication rather than transects, would help to resolve this question.

When considering potential conservation benefits of DFMPAs, it is essential to consider other human uses, most importantly the underlying reason for DFMPA establishment. Unlike MPAs, DFMPAs are generally not managed to achieve conservation goals. Therefore, DFMPAs may have neutral or even negative effects on marine communities, depending on the type and amount of human activity conducted within their boundaries. However, it is unlikely that military activity conducted inside this particular DFMPA has directly adverse effects on marine life. Environmental impact studies conducted at SCI have consistently found that the Navy’s training and testing activities have negligible impact on marine species and habitats [51–52]. Moreover, due to the fact that fishing places such substantial pressure on marine ecosystems, any effects of military activity inside the DFMPA are likely to be substantially less important from a conservation perspective than the associated reduction of fishing pressure.

Understanding the contribution of DFMPAs to marine conservation is critical in the broader context of MSP and current global conservation objectives. In 2010, the United Nations Convention on Biological Diversity (CBD) adopted the Aichi Biodiversity Targets to “safeguard ecosystems, species, and genetic diversity,” among other goals. Aichi Target 11 calls for the protection of at least 17% of terrestrial and inland water areas, and 10% of coastal and marine ecosystems, by 2020. However, many in the scientific and NGO communities are now advocating for an even more aggressive goal – 30% of the world’s ocean protected by 2030. With only 3.5% of the ocean currently protected by MPAs, protection will have to steeply increase for these goals to be met. However, Target 11 also includes “other effective area-based conservation measures” (OECMs) as a potential alternative to formal protected areas for meeting spatial protection goals. In 2018, the Conference of the Parties to the CBD adopted the following definition of an OECM:

> *“A geographically defined area other than a Protected Area, which is governed and managed in ways that achieve positive and sustained long-term outcomes for the in situ conservation of biodiversity with associated ecosystem functions and services and where applicable, cultural, spiritual, socio–economic, and other locally relevant values.”* [53].

Many DFMPAs could fit this definition, illustrating the need to better consider these unique areas and their potential conservation benefits in tracking global progress toward protection goals. Furthermore, given the urgent need for collaborative partnerships in marine conservation, site managers may be interested in working with scientific, conservation, and indigenous communities to achieve both the primary management goal of the DFMPA (e.g. military priorities) as well as biodiversity conservation. More robust monitoring and assessment of DFMPAs, and a better understanding of their potential conservation benefits, are needed before this case can be compellingly made.

To our knowledge, this study is the first spatially explicit, community-wide comparison of marine ecosystems inside and outside a DFMPA. It provides early evidence that DFMPAs may provide conservation benefits similar to those of MPAs. Our results encourage further exploration of the role that DFMPAs may play in marine conservation, and especially their potential integration into existing MSP frameworks and plans to achieve global conservation goals.

## Acknowledgments

Generous financial support for this project came from the United States Pacific Fleet (US Army Corps of Engineers Award W9126G-12-2-0041) and private donations. We thank the crews of the F/V *Donna Kathleen* and Marine Applied Research and Exploration for key assistance in the field.

## Supporting information

**Table S1. Length-weight relationship parameter values and sources for focal fish species.**

**Table S2. Variables (name, year, site, percent rock, mean depth, and area surveyed) associated with ROV transects.**

**Table S3. Means and standard errors for focal species density at fished and DFMPA sites.**

**Table S4. Means and standard errors for focal species biomass at fished and DFMPA sites.**

